# Epigenetic scores for the circulating proteome as tools for disease prediction

**DOI:** 10.1101/2020.12.01.404681

**Authors:** Danni A Gadd, Robert F Hillary, Daniel L McCartney, Shaza B Zaghlool, Anna J Stevenson, Cliff Nangle, Archie Campbell, Robin Flaig, Sarah E Harris, Rosie M Walker, Liu Shi, Elliot M Tucker-Drob, Christian Gieger, Annette Peters, Melanie Waldenberger, Johannes Graumann, Allan F McRae, Ian J Deary, David J Porteous, Caroline Hayward, Peter M Visscher, Simon R Cox, Kathryn L Evans, Andrew M McIntosh, Karsten Suhre, Riccardo E Marioni

## Abstract

Protein biomarkers have been identified across many age-related morbidities. However, characterising epigenetic influences could further inform disease predictions. Here, we leverage epigenome-wide data to study links between the DNAm signatures of the circulating proteome and incident diseases. Using data from four cohorts, we trained and tested epigenetic scores (EpiScores) for 953 plasma proteins, identifying 109 scores that explained between 1% and 58% of the variance in protein levels after adjusting for known protein quantitative trait loci (pQTL) genetic effects. By projecting these EpiScores into an independent sample, (Generation Scotland; n=9,537) and relating them to incident morbidities over a follow-up of 14 years, we uncovered 137 EpiScore – disease associations. These associations were largely independent of immune cell proportions, common lifestyle and health factors and biological aging. Notably, we found that our diabetes-associated EpiScores highlighted previous top biomarker associations from proteome-wide assessments of diabetes. These EpiScores for protein levels can therefore be a valuable resource for disease prediction and risk stratification.

## Introduction

Chronic morbidities place longstanding burdens on our health as we age. Stratifying an individual’s risk prior to symptom presentation is therefore critical (NHS England, 2016). Though complex morbidities are partially driven by genetic factors (Fuchsberger et al., 2016; Yao et al., 2018), epigenetic modifications have also been associated with disease (Lord & Cruchaga, 2014). DNA methylation (DNAm) encodes information on the epigenetic landscape of an individual and blood-based DNAm signatures have been found to predict all-cause mortality and disease onset, providing strong evidence to suggest that methylation is an important measure of disease risk (Hillary, Stevenson, et al., 2020; Lu et al., 2019; Y. Zhang et al., 2017). DNAm can regulate gene transcription (Lea et al., 2018), and epigenetic differences can be reflected in the variability of the proteome (Hillary et al., 2019; Hillary, Trejo-Banos, et al., 2020; Zaghlool et al., 2020). Low-grade inflammation, which is thought to exacerbate many age-related morbidities, is particularly well-captured through DNAm studies of plasma protein levels (Zaghlool et al., 2020). Connecting the epigenome, proteome and time to disease onset may help to identify predictive biological signatures.

Epigenetic predictors have utilised DNAm from the blood to estimate a person’s ‘biological age’ (Lu et al., 2019), measure their exposure to lifestyle and environmental exposures (McCartney, Hillary, et al., 2018; McCartney, Stevenson, Hillary, et al., 2018; Peters et al., 2021) and predict circulating levels of inflammatory proteins (A. Stevenson et al., 2020; A. J. Stevenson et al., 2021). A leading epigenetic predictor of biological aging, the GrimAge epigenetic clock incorporates methylation scores for seven proteins along with smoking and chronological age, and is associated with numerous incident disease outcomes (Hillary, Stevenson, et al., 2020; Lu et al., 2019). This suggests that there is predictive value in utilising DNAm relevant to protein levels for disease predictions. A portfolio of protein EpiScores across the circulating proteome may aid in the prediction of disease and may offer a complementary signal to that of composite scores. Generation of an extensive range of proteomic scores has not been attempted to date. The capability of specific protein scores to predict a range of morbidities has also not been tested. However, DNAm scores for Interleukin-6 and C-Reactive protein have been found to associate with a range of phenotypes independently of measured protein levels, show more stable longitudinal trajectories than repeated protein measurements, and, in some cases, outperform blood-based proteomic associations with brain morphology (Conole et al., 2020; A. J. Stevenson et al., 2021). This is likely due to DNA methylation reflecting a more consistent profile of stress in the body than protein measurements.

Here, we report a comprehensive association study of blood-based DNAm with proteomics and disease (**Figure 1**). We trained epigenetic scores – referred to as EpiScores – for 953 plasma proteins (with sample size ranging from 725 – 944 individuals) and validated them using two independent cohorts with 778 and 162 participants. We regressed out known genetic pQTL effects from the protein levels prior to generating the EpiScores to preclude the signatures being driven by common SNP data that are invariant across the lifespan. Finally, we examined whether the most robust predictors (n=109 EpiScores) associated with the incidence of 12 major morbidities (**Table 1**), over a follow up period of up to 14 years in the Generation Scotland cohort (n = 9,537). We regressed out the effects of age on protein levels prior to training and testing; age was also included as a covariate in the time-to-event disease prediction models. We controlled for common risk factors for disease and assessed the capacity of EpiScores to identify previously reported protein-disease associations.

**Figure 1.**
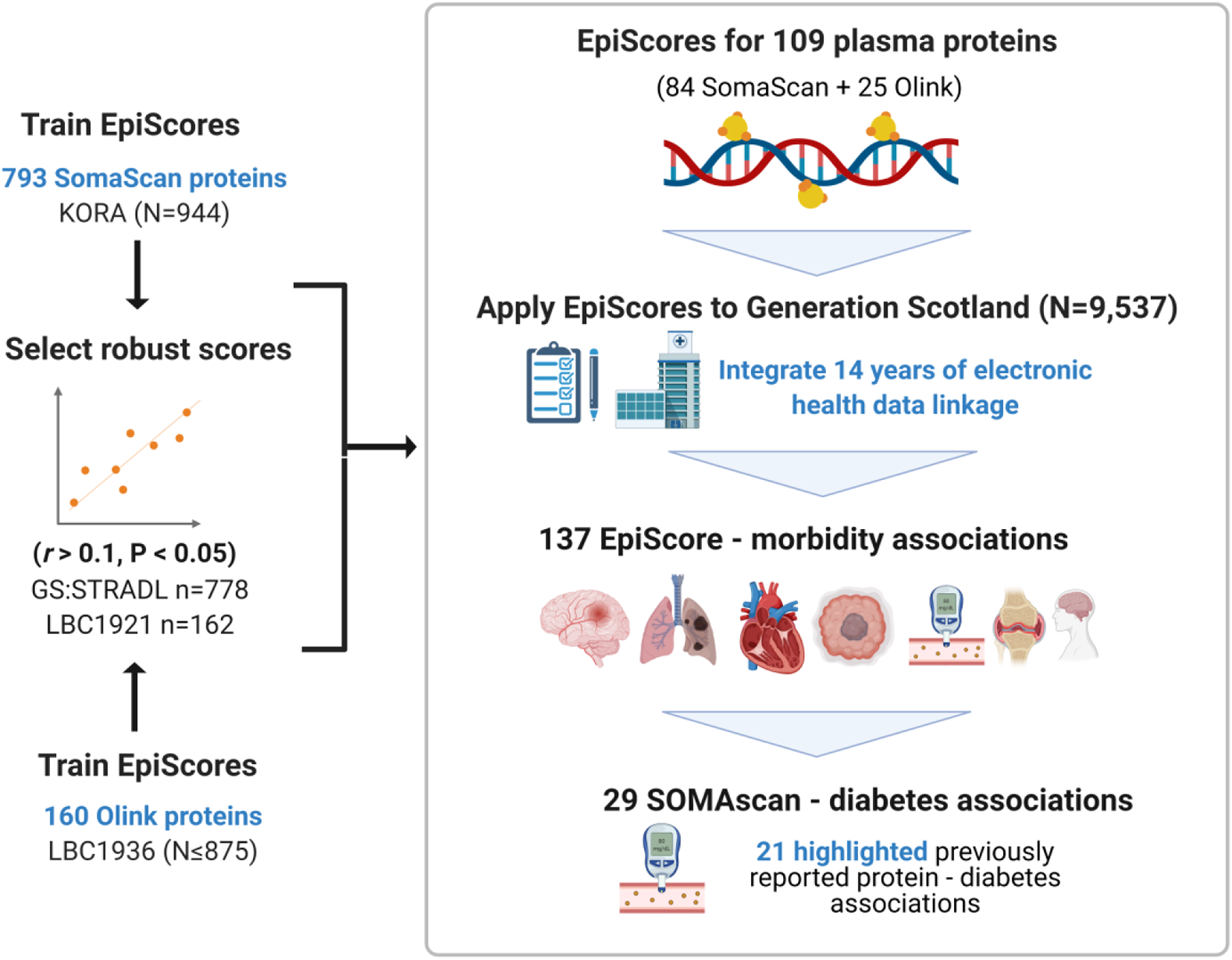
EpiScores for plasma proteins as tools for disease prediction study design. DNA methylation scores were trained on 953 circulating plasma protein levels in the KORA and LBC1936 cohorts. There were 109 EpiScores selected based on performance (*r* > 0.1, P < 0.05) in independent test sets. The selected EpiScores were projected into Generation Scotland, a cohort that has extensive data linkage to GP and hospital records. We tested whether levels of each EpiScore at baseline could predict the onset of 12 leading causes of morbidity, over a follow-up period of up to 14 years. 137 EpiScore – disease associations were identified, for 11 morbidities. We then assessed whether EpiScore associations reflected protein associations for diabetes, which is a trait that has been well-characterised using SOMAscan protein measurements. Of the 29 SOMAscan-derived EpiScore – diabetes associations, 21 reflected highlighted previously reported protein - diabetes associations.

**Table 1.** Incident morbidities in the Generation Scotland cohort. . Counts are provided for the number of cases and controls for each incident trait in the basic and fully-adjusted Cox models run in the Generation Scotland cohort (n=9,537). Mean time-to-event is summarised in years for each phenotype. Alzheimer’s dementia cases and controls were restricted to those older than 65 years. Breast cancer cases and controls were restricted to females.

| Morbidity | Basic model |  |  | Fully-adjusted model |  |  |
| --- | --- | --- | --- | --- | --- | --- |
|  | N cases | N Controls | Years to event (mean, sd) | N cases | N Controls | Years to event (mean, sd) |
| Rheumatoid arthritis | 65 | 9281 | 6.1 (3.5) | 54 | 7736 | 6.4 (3.3) |
| Alzheimer's dementia | 69 | 3764 | 8.3 (2.7) | 52 | 3137 | 8.2 (2.7) |
| Bowel cancer | 77 | 9398 | 6.4 (3.2) | 65 | 7817 | 6.5 (3.2) |
| Depression | 101 | 8306 | 3.9 (3.3) | 80 | 6976 | 3.7 (3.3) |
| Breast cancer | 129 | 5355 | 6 (3.4) | 110 | 4401 | 5.9 (3.4) |
| Lung cancer | 201 | 9265 | 5.2 (3.1) | 172 | 7705 | 5.1 (3.1) |
| Inflammatory bowel disease | 203 | 9083 | 5 (3.5) | 163 | 7567 | 4.9 (3.5) |
| Stroke | 317 | 9023 | 6.5 (3.4) | 248 | 7546 | 6.4 (3.5) |
| COPD | 346 | 8939 | 6.2 (3.4) | 273 | 7459 | 6.1 (3.4) |
| Ischaemic heart disease | 395 | 8646 | 5.8 (3.3) | 309 | 7248 | 5.9 (3.3) |
| Diabetes | 428 | 8756 | 5.7 (3.4) | 322 | 7331 | 5.7 (3.4) |
| Pain | 1494 | 5341 | 5.2 (3.5) | 1221 | 4475 | 5.3 (3.5) |

## Results

### Selecting the most robust EpiScores for protein levels

To generate epigenetic scores for a comprehensive set of plasma proteins, we ran elastic net penalised regression models using protein measurements from the SOMAscan (aptamer-based) and Olink (antibody-based) platforms. We used two cohorts: the German population-based study KORA (n=944, mean age 59 years (SD 7.8), with 793 SOMAscan proteins) and the Scottish Lothian Birth Cohort 1936 (LBC1936) study (between 725 and 875 individuals in the training cohort, with a total of 160 Olink neurology and inflammatory panel proteins). The mean age of the LBC1936 participants at sampling was 70 (SD 0.8) for inflammatory and 73 (SD 0.7) for neurology proteins. Full demographic information is available for all cohorts in **Supplementary file 1A**.

Prior to running the elastic net models, we rank-based inverse normalised protein levels and adjusted for age, sex, cohort-specific variables and, where present, *cis* and *trans* pQTL effects identified from previous analyses (Hillary et al., 2019; Hillary, Trejo-Banos, et al., 2020; Suhre et al., 2017) (**Methods**). Of a possible 793 proteins in KORA, 84 EpiScores had Pearson *r* > 0.1 and P < 0.05 when tested in an independent subset of Generation Scotland (The Stratifying Resilience and Depression Longitudinally [STRADL] study, n=778) (**Supplementary file 1B**). These EpiScores were selected for EpiScore-disease analyses. Of the 160 Olink proteins trained in LBC1936, there were 21 with *r* > 0.1 and P < 0.05 in independent test sets (STRADL, n=778, Lothian Birth Cohort 1921: LBC1921, n=162) (**Supplementary file 1C**). Independent test set data were not available for four Olink proteins. However, they were included based on their performance (*r* > 0.1 and P < 0.05) in a holdout sample of 150 individuals who were left out of the training set. We then retrained these four predictors on the full training sample.

A total of 109 EpiScores (84 SOMAscan-based and 25 Olink-based) were brought forward (*r* > 0.1 and P < 0.05) to EpiScore-disease analyses (**Figure 2 and Supplementary file 1D**). There were five EpiScores for proteins common to both Olink and SOMAscan panels, which had variable correlation strength (GZMA *r* = 0.71, MMP.1 *r* = 0.46, CXCL10 *r* = 0.35, NTRK3 *r* = 0.26, and CXCL11 *r* = 0.09). Predictor weights, positional information and *cis/trans* status for CpG sites contributing to these EpiScores are available in **Supplementary file 1E**. The number of CpG features selected for EpiScores ranged from one (Lyzozyme) to 395 (Aminoacylase-1), with a mean of 96 **Supplementary file 1F**). The most frequently selected CpG was the smoking-related site cg05575921 (mapping to the *AHRR* gene), which was included in 25 EpiScores. Counts for each CpG site are summarised in **Supplementary file 1G.** This table includes the set of protein EpiScores that each CpG contributes to, along with phenotypic annotations (traits) from the MRC-IEU EWAS catalog (MRC-IEU, 2021) for each CpG site having genome-wide significance (P < 3.6 x10^-8^) (Saffari et al., 2017).

**Figure 2.**
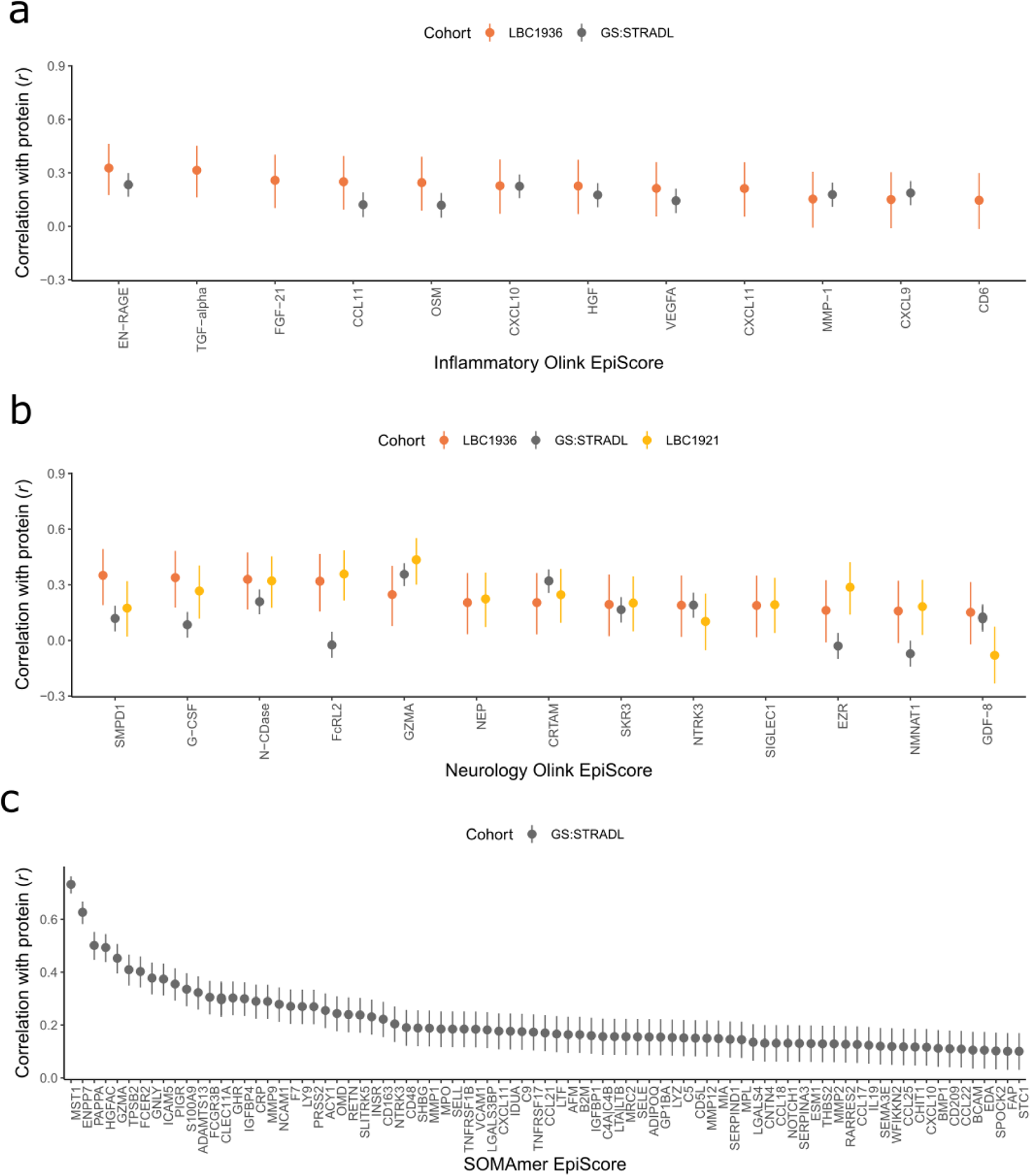
Test performance for the 109 selected protein EpiScores. Test set correlation coefficients for associations between protein EpiScores for (**a**) inflammatory Olink, (**b**) neurology Olink and (**c**) SOMAmer protein panel EpiScores and measured protein levels are plotted. Upper and lower confidence intervals are shown for each correlation. The 109 protein EpiScores shown achieved *r* > 0.1 and P < 0.05 either one or both of the GS:STRADL (n=778) and LBC1921 (n=162) test sets, wherever protein data was available for comparison. Data shown corresponds to the results included in **Supplementary files 1B-C**.

### EpiScore-disease associations in Generation Scotland

The Generation Scotland dataset contains extensive electronic health data from GP and hospital records available as well as DNA methylation data for 9,537 individuals. This makes it uniquely positioned to test whether EpiScore signals can predict disease onset. We ran nested mixed effects Cox proportional hazards models (**Figure 3**) to determine whether the levels of each EpiScore at baseline associated with the incidence of 12 morbidities over a maximum of 14 years of follow up. The correlation structures for the 109 EpiScore measures used for Cox modelling are presented in **Supplementary file 2A**.

**Figure 3.**
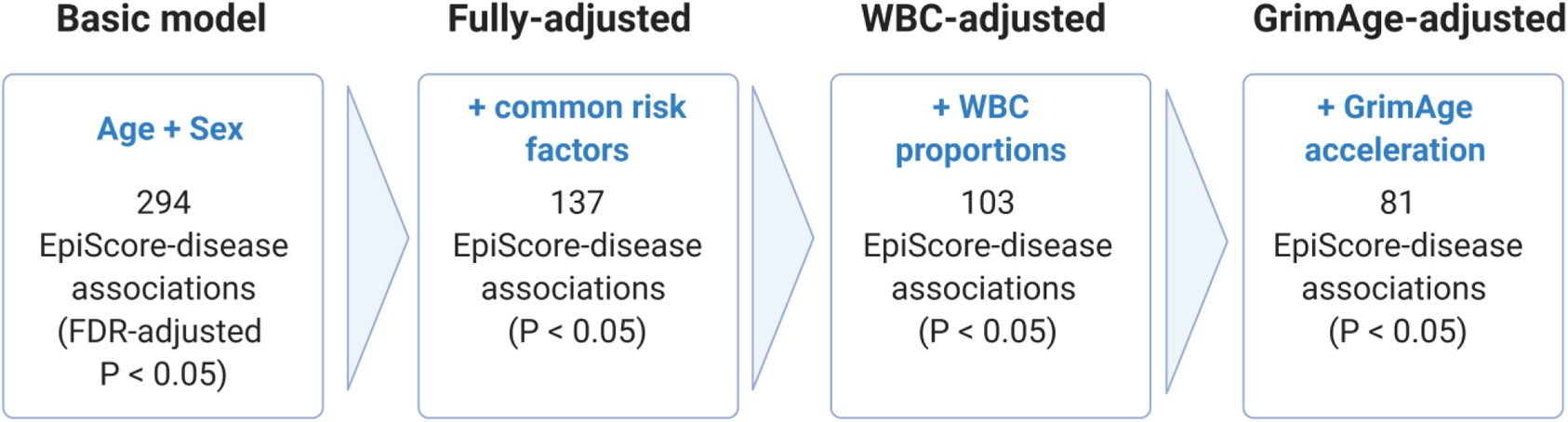
Nested Cox proportional hazards assessment of EpiScore-disease prediction. Mixed effects Cox proportional hazards analyses in Generation Scotland (n = 9,537) tested the relationships between each of the 109 selected EpiScores and the incidence of 12 leading causes of morbidity **(Supplementary files 1H-I)**. The basic model was adjusted for age and sex and yielded 294 associations between EpiScores and disease diagnoses, with FDR-adjusted P < 0.05. In the fully-adjusted model, which included common risk factors as additional covariates (smoking, deprivation, educational attainment, BMI and alcohol consumption) 137 of the basic model associations remained significant with P < 0.05. In a sensitivity analysis, the addition of estimated White Blood Cells (WBCs) to the fully-adjusted models led to the attenuation of 34 of the 137 associations. In a further sensitivity analysis, 81 associations remained after adjustment for both immune cell proportions and GrimAge acceleration.

The Cox proportional hazard assumption dictates that hazard ratios for EpiScore – disease associations should remain constant over time. We correlated the Schoenfeld residuals from the models with time to test this. Two associations in the basic model adjusting for age and sex failed to satisfy the global assumption (across all covariates) and were excluded. There were 294 remaining EpiScore-disease associations with a False Discovery Rate (FDR)-adjusted P < 0.05 in the basic model. After further adjustment for common risk factor covariates (smoking, social deprivation status, educational attainment, body mass index (BMI) and alcohol consumption), 137 of the 294 EpiScore-disease associations from the basic model had P < 0.05 in the fully-adjusted model (**Supplementary files 1H-I**). Eleven of the 137 fully-adjusted associations failed the Cox proportional hazards assumption for the EpiScore variable (P < 0.05 for the association between the Schoenfeld residuals and time; **Supplementary file 1J**). When we restricted the time-to-event/censor period by each year of possible follow-up, there were minimal differences in the EpiScore - disease hazard ratios between follow-up periods that did not violate the assumption and those that did (**Supplementary file 1K**). The 137 associations were therefore retained as the primary results.

The 137 associations found in the fully-adjusted model comprised 78 unique EpiScores that were related to the incidence of 11 of the 12 morbidities studied. Diabetes and chronic obstructive pulmonary disease (COPD) had the greatest number of associations, with 33 and 41, respectively. **Figure 4** presents the EpiScore-disease relationships for COPD and the remaining nine morbidities: stroke, lung cancer, ischaemic heart disease, inflammatory bowel disease, rheumatoid arthritis, depression, bowel cancer, pain and Alzheimer’s dementia. There were 13 EpiScores that associated with the onset of three or more morbidities. **Figure 5** presents relationships for these 13 EpiScores in the fully-adjusted Cox model results. Of note is the EpiScore for Complement 5 (C5), which associated with five outcomes: stroke, diabetes, ischaemic heart disease, rheumatoid arthritis and COPD. Of the 29 SOMAscan-derived EpiScore associations with incident diabetes, 21 replicated previously reported protein associations (Elhadad et al., 2020; Gudmundsdottir et al., 2020) with incident or prevalent diabetes in one or more cohorts (**Figure 6 and Supplementary file 1L**).

**Figure 4.**
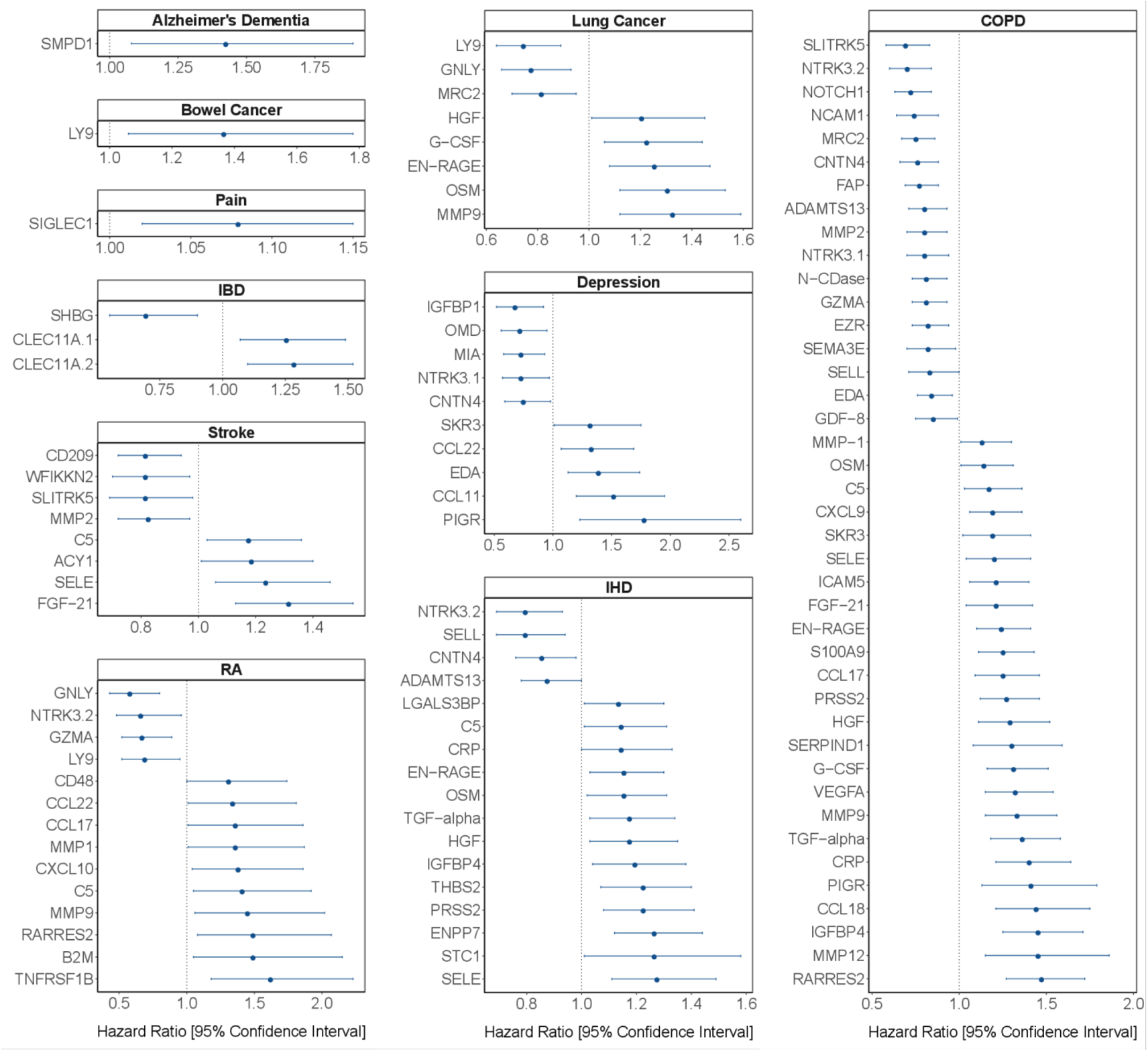
EpiScore associations with incident disease. EpiScore-disease associations for ten of the eleven morbidities with associations where P < 0.05 in the fully-adjusted mixed effects Cox proportional hazards models in Generation Scotland (n=9,537). Hazard ratios are presented with confidence intervals for 104 of the 137 EpiScore – incident disease associations reported. Models were adjusted for age, sex and common risk factors (smoking, BMI, alcohol consumption, deprivation and educational attainment). IBD: inflammatory bowel disease. IHD: ischaemic heart disease. COPD: chronic obstructive pulmonary disease. For EpiScore - diabetes associations, see Figure 6. Data shown corresponds to the results included in **Supplementary file 1I**.

**Figure 5.**
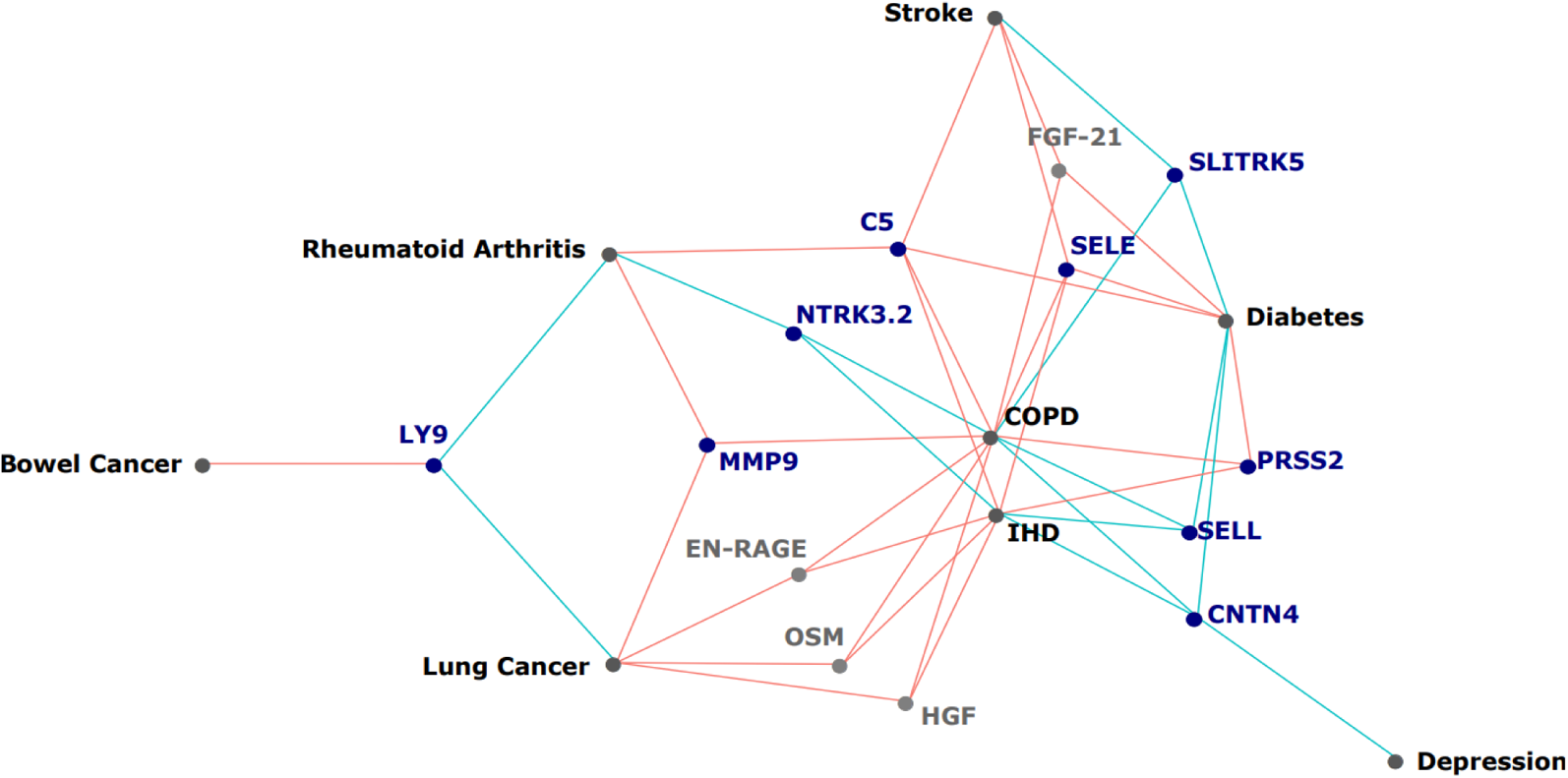
EpiScores that associated with the greatest number of morbidities. EpiScores with a minimum of three relationships with incident morbidities in the fully-adjusted Cox models. The network includes 13 EpiScores as dark blue (SOMAscan) and grey (Olink) nodes, with disease outcomes in black. EpiScore-disease associations with hazard ratios < 1 are shown as blue connections, whereas hazard ratios > 1 are shown in red. COPD: chronic obstructive pulmonary disease. IHD: ischaemic heart disease. Data shown corresponds to the results included in **Supplementary file 1I**.

**Figure 6.**
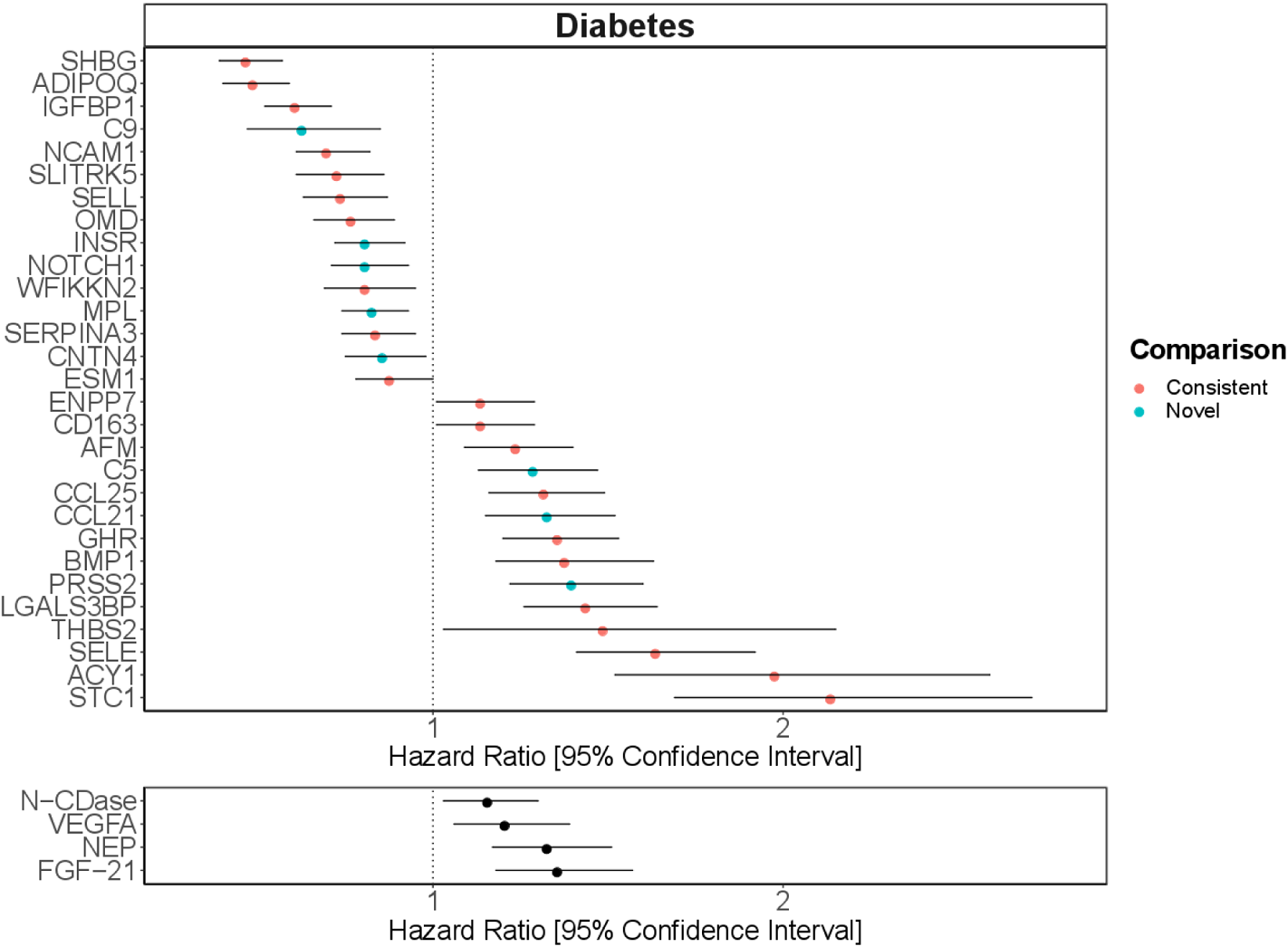
Replication of known protein-diabetes associations with EpiScores. EpiScore – incident diabetes associations in Generation Scotland (n=9,537). The 29 SOMAscan (top panel) and four Olink (bottom panel) associations shown with P < 0.05 in fully-adjusted mixed effects Cox proportional hazards models. Of the 29 SOMAscan-derived EpiScores, 21 associations were consistent with protein – diabetes associations (pink) in one or more of the four comparison cohorts that used SOMAscan protein levels. Eight associations were novel (blue). Data shown corresponds to the results included in **Supplementary files 1I and 1L** .

### Immune cell and GrimAge sensitivity analyses

Correlations of the 109 EpiScores with covariates suggested interlinked relationships with both immune cells and GrimAge acceleration (**Supplementary file 2B)**. These covariates were therefore added incrementally to the fully-adjusted Cox models (**Figure 3**). There were 103 associations that remained statistically significant (FDR P < 0.05 in the basic model and P < 0.05 in the fully-adjusted model) after adjustment for immune cell proportions, of which 81 remained significant when GrimAge acceleration scores were added to this model **(Supplementary file 1I)**. In a further sensitivity analysis, relationships between both estimated White Blood Cell (WBC) proportions and GrimAge acceleration scores with incident diseases were assessed in the Cox model structure independently of EpiScores. Of the 60 possible relationships between WBC measures and the morbidities assessed, four were statistically significant (FDR-adjusted P < 0.05) in the basic model and remained significant with P < 0.05 in the fully-adjusted model **(Supplementary file 1M**). A higher proportion of Natural Killer cells was linked to decreased risk of incident COPD, rheumatoid arthritis, diabetes and pain. The GrimAge acceleration composite score was associated with COPD, IHD, Diabetes and Pain in the fully-adjusted models (P < 0.05) (**Supplementary file 1N**). The magnitude of the GrimAge effect sizes were comparable to the EpiScore findings.

## Discussion

Here, we report a comprehensive DNA methylation scoring study of 953 circulating proteins. We define 109 robust EpiScores for plasma protein levels that are independent of known pQTL effects. By projecting these EpiScores into a large cohort with extant data linkage, we show that 78 EpiScores associate with the incidence of 11 leading causes of morbidity (137 EpiScore disease associations in total). Finally, we show that EpiScore - disease associations highlight previously measured protein - disease relationships. The bulk of EpiScore-disease associations are independent of common lifestyle and health factors, differences in immune cell composition and GrimAge acceleration. EpiScores therefore provide methylation-proteomic signatures for disease prediction and risk stratification.

The consistency between our EpiScore – diabetes associations and previously identified protein – diabetes relationships (Elhadad et al., 2020; Gudmundsdottir et al., 2020) suggests that epigenetic scores may identify candidate disease-protein pathways. In addition to the comprehensive lookup of SOMAscan proteins with diabetes, several of the markers we identified for COPD and IHD also reflect previous associations with measured proteins (Ganz et al., 2016; Serban et al., 2021). The two studies used for the diabetes comparison represent the largest candidate protein characterisations of diabetes to date and the top markers identified included aminoacylase-1 (ACY-1), sex hormone binding globulin (SHBG), growth hormone receptor (GHR) and Insulin-like growth factor-binding protein 2 (IGFBP-2) (Elhadad et al., 2020; Gudmundsdottir et al., 2020). Our EpiScores for these top markers are also associated with diabetes, in addition to EpiScores for several other protein markers reported in these studies. A growing body of evidence suggests that type 2 diabetes is mediated by genetic and epigenetic regulators (Kwak & Park, 2016) and proteins such as ACY-1 and GHR are thought to influence a range of diabetes-associated metabolic mechanisms (Kim & Park, 2017; Pérez-Pérez et al., 2012). In the case of diabetes, EpiScores may therefore be used as disease-relevant risk biomarkers, many years prior to onset. Validation should be tested when sufficient data become available for the remaining morbidities.

With modest test set performances (for example, SHBG *r* = 0.18 and ACY-1 *r* = 0.25), it is perhaps surprising that such strong synergy is observed between EpiScores for proteins that associated with diabetes and the trends seen with measured proteins. Nonetheless, DNA methylation scores for CRP and IL6 have been shown to perform modestly in test sets (*r* ∼ 0.2, equivalent to ∼ 4% explained variance in protein level), but augment and often outperform the measured protein related to a range of phenotypes (A. Stevenson et al., 2020; A. J. Stevenson et al., 2021). Upper bounds for DNAm prediction of complex traits, such as proteins, can be estimated by variance components analyses (Hillary, Trejo-Banos, et al., 2020; Trejo Banos et al., 2020; F. Zhang et al., 2019).

Compared to epigenetic clocks like GrimAge, EpiScores enable the granular study of individual protein predictor signatures with disease outcomes. For example, levels of the acid sphingomyelinase (ASM) EpiScore predicted onset of Alzheimer’s dementia, several years prior to diagnosis. ASM (encoded by *SMPD1*) has been discussed as a therapeutic candidate for Alzheimer’s disease (Cataldo et al., 2004; Kamil et al., 2016; Lee et al., 2014; Park et al., 2020) and has been shown to disrupt autophagic protein degradation and associate with accumulation of amyloid-beta in murine models of Alzheimer’s pathology (Lee et al., 2014; Park et al., 2020). The EpiScore for Complement Component 5 (C5) was associated with the onset of five morbidities, the highest number for any EpiScore. Elevated levels of C5 peptides have been associated with severe inflammatory, autoimmune and neurodegenerative states (Ma et al., 2019; Mantovani et al., 2014; Morgan & Harris, 2015) and a range of C5-targetting therapeutic approaches are in development (Alawieh et al., 2018; Brandolini et al., 2019; Hawksworth et al., 2017; Hernandez et al., 2017; Morgan & Harris, 2015; Ort et al., 2020). EpiScores that occupy central hubs in the disease-prediction framework may therefore provide evidence of early methylation signatures common to the onset of multiple diseases. Our large-scale assessment of EpiScores provides a platform for future studies, as composite predictors may be created using our EpiScore database. These should be tested in incident disease predictions when sufficient case data are available.

This study has several limitations. First, like with protein – disease association studies, we cannot infer causality from our EpiScore – disease models. However, both protein levels and EpiScores may have utility in risk prediction – future studies where both modalities are available should assess paired protein and CpG contributions to traits. This should entail the direct measurement of proteins, as inference from EpiScores alone, while useful for disease risk stratification, is not sufficient to determine mechanisms. Second, the epitope nature of the protein measurement in the SOMAscan panel may incur probe cross-reactivity and non-specific binding; there may also be differences in how certain proteins are measured across panels (Pietzner et al., 2020; Sun et al., 2018). Comparisons of both protein measurement technologies on the same samples would help to explore this in more detail. Third, there may also be pQTLs with small effect sizes that were not regressed from the proteins prior to generating the EpiScores. Finally, associations present between EpiScore measures and disease incidence may have been influenced by external factors such as prescription medications for comorbid conditions and comorbid disease prevalence.

We have shown that EpiScores for circulating protein levels predict the incidence of multiple diseases, up to 14 years prior to diagnosis. Our findings suggest that DNA methylation phenotyping approaches and data linkage to electronic health records in large, population-based studies have the potential to (1) Capture inter-individual variability in protein levels; (2) Augment risk prediction many years prior to morbidity onset; and (3) highlight candidate protein – disease associations for further exploration. The EpiScore weights are publicly available, enabling any cohort with Illumina DNAm data to generate them and to relate them to various outcomes. Given the increasingly widespread assessment of DNAm in cohort studies (McCartney et al., 2020; Min et al., 2020), EpiScores offer an affordable and consistent (i.e. array-based) way to utilise these signatures. This information is likely to be important in risk stratification and prevention of age-related morbidities.

## Materials and Methods

### The KORA sample population

The KORA F4 study includes 3,080 participants who reside in Southern Germany. Individuals were between 32 and 81 years of age when recruited to the study from 2006 and 2008. In the current study, there were 944 individuals with methylation, proteomics and genetic data available. The Infinium HumanMethylation450 BeadChip platform was used to generate DNA methylation data for these individuals. The Affymetrix Axiom array was used to generate genotyping data and the SOMAscan platform was used to generate proteomic measurements in the sample.

### DNA methylation in KORA

Methylation data were generated for 1,814 individuals(Petersen et al., 2014); 944 also had protein and genotype measurements available. During preprocessing, 65 SNP probes were excluded and background correction was performed in minfi (Aryee et al., 2014). Samples with a detection rate of less than 95% were excluded. Next, the minfi R package was used to perform normalization on the intensity methylation measures (Aryee et al., 2014), with a method consistent with the Lumi:QN +BMIQ pipeline. After excluding non-cg sites and CpGs on sex chromosomes or with fewer than 100 measures available, 470,837 CpGs were available for analyses.

### Proteomics in KORA

The SOMAscan platform (V3.2) (Gold et al., 2010) was used to quantify protein levels in undepleted plasma for 1129 SOMAmer probes (Suhre et al., 2017). Of the 1,000 samples provided for analysis, two samples were excluded due to errors in bio-bank sampling and one based on quality control measures. Of the 997 samples available, there were 944 individuals with methylation and genotypic data. Of the 1,129 probes available, five failed the QC, leaving a total of 1,124 probes for the subsequent analysis. Protein measurements were transformed by rank-based inverse normalisation and regressed onto age, sex, known pQTLs and 20 genetic principal components of ancestry derived from the Affymetrix Axiom Array to control for population structure. pQTLs for each protein were taken from a previous GWAS in the sample (Suhre et al., 2017).

### The LBC1936 and LBC1921 sample populations

The Lothian Birth Cohorts of 1921 (LBC1921; N = 550) and 1936 (LBC1936; N = 1091) are longitudinal studies of aging in individuals who reside in Scotland (Deary et al., 2012; Taylor et al., 2018). Participants completed an intelligence test at age 11 years and were recruited for these cohorts at mean ages of 79 (LBC1921) and 70 (LBC1936). They have been followed up triennially for a series of cognitive, clinical, physical and social data, along with blood donations that have been used for genetic, epigenetic, and proteomic measurement. DNAm, proteomic (Olink® platform) and genetic data for up to 875 individuals from Waves 1 and 2 of the LBC1936 (at mean ages 70 and 73 years) and 162 individuals at Wave 3 of the LBC1921 (at mean age 87 years).

### DNAm in LBC1936 and LBC1921

DNA from whole blood was assessed using the Illumina 450 K methylation array. Details of quality control have been described elsewhere (Shah et al., 2014; Q. Zhang et al., 2018). Raw intensity data were background-corrected and normalised using internal controls. Manual inspection resulted in the removal of low quality samples that presented issues related to bisulphite conversion, staining signal, inadequate hybridisation or nucleotide extension. Probes with low detection rate <95% at P < 0.01 and samples with low call rates (<450,000 probes detected at P < 0.01) were removed. Samples were also removed if they had a poor match between genotype and SNP control probes, or incorrect DNA methylation-predicted sex.

### Proteomics in LBC1936 and LBC1921

Plasma samples were analysed using either the Olink® neurology 92-plex or the Olink® inflammation 92-plex proximity extension assays (Olink® Bioscience, Uppsala Sweden). One inflammatory panel protein (BDNF) failed quality control and was removed. A further 21 proteins were removed, as over 40% of samples fell below the lowest limit of detection. Two neurology proteins, MAPT and HAGH, were excluded due to >40% of observations being below the lower limit of detection. This resulted in 90 neurology (LBC1936 Wave 2) and 70 inflammatory (LBC1936 Wave 1) proteins in LBC1936 and 92 neurology proteins available in LBC1921. Protein levels were rank-based inverse normalised and regressed onto age, sex, four genetic components of ancestry derived from multidimensional scaling of the Illumina 610-Quadv1 genotype array and Olink® array plate. In LBC1936, pQTLs were adjusted for, through reference to GWAS in the samples (Hillary et al., 2019; Hillary, Trejo-Banos, et al., 2020).

### Generation Scotland and STRADL sample populations

Generation Scotland: the Scottish Family Health Study (GS) is a large, family-structured, population-based cohort study of >24,000 individuals from Scotland (mean age 48 years) (Smith et al., 2006). Recruitment took place between 2006 and 2011 with a clinical visit where detailed health, cognitive, and lifestyle information was collected along with biological samples (blood, urine, saliva). In GS, there were 9,537 individuals with DNAm and phenotypic information available. The Stratifying Resilience and Depression Longitudinally (STRADL) cohort is a subset of 1,188 individuals from the GS cohort who undertook additional assessments approximately five years after the study baseline (Navrady et al., 2018).

### DNA methylation in Generation Scotland and STRADL

In the GS cohort, blood-based DNA methylation was generated in two sets using the Illumina EPIC array. Set 1 comprised 5,190 related individuals whereas Set 2 comprised 4,583 individuals, unrelated to each other and to those in Set 1. During quality control, probes were removed based on visual outlier inspection, bead count <3 in over 5% of samples and samples with detection P value below adequate thresholds (McCartney, Stevenson, Walker, et al., 2018; Seeboth et al., 2020). Samples were removed based on sex mismatches, low detection P values for CpGs and saliva samples and genetic outliers (Amador et al., 2015). The quality-controlled dataset comprised 9,537 individuals (n_Set1_=5,087, n_Set2_=4,450). The same steps were also applied to process DNAm in STRADL.

### Proteomics in STRADL

Measurements for 4,235 proteins in 1,065 individuals from the STRADL cohort were recorded using the SOMAscan® technology. 793 epitopes matched between the KORA and STRADL cohorts and were included for training in KORA and testing in STRADL. There were 778 individuals with proteomics data and DNAm data in STRADL. Protein measurements were transformed by rank-based inverse normalisation and regressed onto age, sex and 20 genetic principal components (derived from multidimensional scaling of genotype data from the Illumina 610-Quadv1 array).

### Electronic health data linkage in Generation Scotland

Over 98% of GS participants consented to allow access to electronic health records via data linkage to GP records (Read 2 codes) and hospital records (ICD codes). Data are available prospectively from the time of blood draw, yielding up to 14 years of linkage. We considered incident data for 12 morbidities (**Supplementary file 3A**). Prevalent cases (ascertained via retrospective ICD and Read 2 codes or self-report from a baseline questionnaire) were excluded. For inflammatory bowel disease (IBD) prevalent cases were excluded based on data linkage alone. Included and excluded terms can be found in **Supplementary files 4A-L**. Alzheimer’s dementia was limited to cases/controls with age of event/censoring ≥ 65 years. Breast cancer was restricted to females only. Recurrent, major and moderate episodes of depression were included in depression. Diabetes was comprised of type 2 diabetes and more general diabetes codes such as diabetic retinopathy and diabetes mellitus with renal manifestation. Type 1 and juvenile diabetes cases were excluded.

### Elastic net protein EpiScores

Penalised regression models were generated for 160 proteins in LBC1936 and 793 proteins in KORA using Glmnet (Version 4.0-2) (J et al., 2010) in R (Version 4.0) (R, 2020). Protein levels were the outcome and there were 428,489 CpG features per model in the LBC1936 training and 397,630 in the KORA training. An elastic net penalty was specified (alpha=0.5) and cross validation was applied. DNAm and protein measurements were scaled to have a mean of zero and variance of one.

In the KORA analyses, 10-fold cross validation was applied and EpiScores were tested in STRADL (n=778). Of 480 EpiScores that generated ≥1 CpG features, 84 had Pearson r > 0.1 and P < 0.05 in STRADL. As test set comparisons were not available for every protein in the LBC1936 analyses, a holdout sample was defined, with two folds set aside as test data and 10-fold cross validation carried out on the remaining data (n_train_=576, n_test_=130 for neurology and n_train_=725, n_test_=150 for inflammatory proteins). We retained 36 EpiScores with Pearson r > 0.1 and P < 0.05. New predictors for these 36 proteins were then generated using 12-fold cross validation and tested externally in STRADL (n=778) and LBC1921 (n=162, for the neurology panel). 21 EpiScores had r > 0.1 and P < 0.05 in at least one of the external test sets. Four EpiScores did not have external comparisons and were included based on holdout performance. The 109 selected EPiScores were then applied to Generation Scotland (n=9,537). DNAm at each CpG site was scaled to have a mean of zero and variance of one, with scaling performed separately for GS Sets.

### Associations with health linkage phenotypes in Generation Scotland

Mixed effects Cox proportional hazards regression models adjusting for age, sex, and methylation set were used to assess the relationship between 109 EpiScores and 12 morbidities in Generation Scotland. Models were run using coxme (Therneau, 2020b) (Version 2.2-16) with a kinship matrix accounting for relatedness in Set 1. Cases included those diagnosed after baseline who had died, in addition to those who received a diagnosis and remained alive. Controls were censored if disease free at time of death, or at the end of the follow-up period. EpiScore levels were rank-base inverse normalised. Fully-adjusted models included: the following additional covariates measured at baseline: alcohol consumption (units consumed in the previous week); deprivation (assessed by the Scottish Index of Multiple Deprivation (GovScot, 2016)); body mass index (kg/m^2^); educational attainment (an 11-category ordinal variable) and a DNAm-based score for smoking status (Bollepalli et al., 2019). A false discovery rate multiple testing correction P < 0.05 was applied to the 1306 EpiScore-disease associations (109 EpiScores by 12 incident disease traits, with 2 associations excluded for failing the global proportional hazards assumption). Proportional hazards assumptions were checked through Schoenfeld residuals (global test and a test for the protein-EpiScore variable) using the coxph and cox.zph functions from the survival package (Therneau, 2020a) (Version 3.2-7). For each association failing to meet the assumption (Schoenfeld residuals P < 0.05), a sensitivity analysis was run across yearly follow-up intervals.

Fully-adjusted Cox proportional hazards models were run with Houseman-estimated White Blood Cell (WBC) proportions as covariates (Houseman et al., 2012). A further sensitivity analyses added GrimAge acceleration (Lu et al., 2019) as an additional covariate. Basic and fully-adjusted Cox models were also run with estimated Monocyte, Bcell, CD4T, CD8T and Natural Killer cell proportions as predictors.

Correlation structures for EpiScores, DNAm-based white cell proportions and phenotypic information were assessed using Pearson correlations and pheatmap (Kolde, 2019) (Version 1.0.12) and ggcorrplot packages (Version 0.1.3) (Kassambara, 2019). The psych package (Version 2.0.9) (Revelle, 2020) was used to perform principal components analysis on EpiScores. A network visualisation was produced using the ggraph package (Version 2.0.5) (Pedersen, 2021). Figures 1 and 2 were created with BioRender.com.

### Consistency of disease associations between EpiScores and measured proteins

Comparisons were conducted between EpiScore – diabetes associations and diabetes associations with measured proteins using two previous large-scale proteomic studies (Elhadad et al., 2020; Gudmundsdottir et al., 2020). In both studies, two cohorts were included (Study 1: KORA n= 993, HUNT n= 940 (Elhadad et al., 2020), Study 2: AGES-Reykjavik n=5,438 and QMDiab n=356 (Gudmundsdottir et al., 2020)). Study 1 included the KORA dataset, which we use in this study to generate SOMAscan EpiScores. We characterised which SOMAscan-based EpiScore – diabetes associations from our fully-adjusted results reflected those observed with measured protein levels. We included basic (nominal P < 0.05) and fully adjusted results (with either FDR or Bonferroni-corrected P < 0.05), wherever available, across the four cohorts (**Supplementary file 1L**).

## Acknowledgements

We are grateful to all study participants of KORA, LBC1936, LBC1921 and GS for their invaluable contributions to this study.

## Funding

**This research was funded in whole, or in part, by the Wellcome Trust [104036/Z/14/Z, 108890/Z/15/Z, 203771/Z/16/Z]. For the purpose of open access, the author has applied a CC BY public copyright licence to any Author Accepted Manuscript version arising from this submission**. The LBC1921 was supported by the UK’s Biotechnology and Biological Sciences Research Council (BBSRC), a Royal Society–Wolfson Research Merit Award to I.J.D., and the Chief Scientist Office (CSO) of the Scottish Government’s Health Directorates. The LBC1936 is supported by Age UK (Disconnected Mind project, which supports S.E.H.), the Medical Research Council (G0701120, G1001245, MR/M013111/1, MR/R024065/1, which supports S.R.C.), and the University of Edinburgh. Genotyping was supported by the Biotechnology and Biological Sciences Research Council (BB/F019394/1). Methylation typing in both the LBC1921 and LBC1936 was supported by Centre for Cognitive Ageing and Cognitive Epidemiology (Pilot Fund award), Age UK, The Wellcome Trust Institutional Strategic Support Fund, The University of Edinburgh, and The University of Queensland. Proteomic analyses in LBC1936 and LBC1921 were supported for by the Age UK grant and NIH Grants R01AG054628 and R01AG05462802S1. This work was conducted in the Centre for Cognitive Ageing and Cognitive Epidemiology, which was supported by the Medical Research Council and Biotechnology and Biological Sciences Research Council (MR/K026992/1) and which supports I.J.D. We acknowledge Grant P2CHD042849 for supporting the Population Research Center at the University of Texas. This research was supported by Australian National Health and Medical Research Council (grants 1010374, 1046880 and 1113400) and by the Australian Research Council (DP160102400). P.M.V. is supported by the NHMRC Fellowship Scheme (1078037, 1078901). A.F.M. is supported by the Australian Research Council Fellowship (FT200100837). P.M.V. was also funded by the Australian Research Council (DP160102400 and FL180100072). Lothian Birth Cohort 1921 and 1936 proteomic analyses were supported by a National Institutes of Health research grant (R01AG054628) which also supports E.M.T-D and S.R.C.

Generation Scotland received core support from the Chief Scientist Office of the Scottish Government Health Directorates (CZD/16/6) and the Scottish Funding Council (HR03006). Genotyping and DNA methylation profiling of the GS samples was carried out by the Genetics Core Laboratory at the Clinical Research Facility, University of Edinburgh, Scotland and was funded by the Medical Research Council UK and the Wellcome Trust (Wellcome Trust Strategic Award “STratifying Resilience and Depression Longitudinally” ([STRADL; Reference 104036/Z/14/Z]). Proteomic analyses in STRADL were supported by Dementias Platform UK (DPUK). DPUK funded this work through core grant support from the Medical Research Council [MR/L023784/2]. C.H. is supported by an MRC University Unit Programme Grant MC_UU_00007/10 (QTL in Health and Disease). L.S. is funded by DPUK through MRC (grant no. MR/L023784/2) and the UK Medical Research Council Award to the University of Oxford (grant no. MC_PC_17215). L.S. also received support from the NIHR Biomedical Research Centre at Oxford Health NHS Foundation Trust.

K.S. and S.B.Z. are supported by the Biomedical Research Program at Weill Cornell Medicine in Qatar, a program funded by the Qatar Foundation. K.S. is also supported by QNRF grant NPRP11C-0115-180010.

The KORA study was initiated and financed by the Helmholtz Zentrum München—German Research Center for Environmental Health, which is funded by the German Federal Ministry of Education and Research (BMBF) and by the State of Bavaria. Furthermore, KORA research was supported within the Munich Center of Health Sciences (MC-Health), Ludwig-Maximilians-Universität, as part of LMUinnovativ. The KORA study is funded by the Bavarian State Ministry of Health and Care through the research project DigiMed Bayern (www.digimed-bayern.de). We gratefully acknowledge the contribution of all field staff conducting the KORA F4 study.

D.A.G., R.F.H. and A.J.S. are supported by funding from the Wellcome Trust 4-year PhD in Translational Neuroscience–training the next generation of basic neuroscientists to embrace clinical research [D.A.G. and R.F.H.: 108890/Z/15/Z; A.J.S: 203771/Z/16/Z]. D.L.Mc.C. and R.E.M. are supported by Alzheimer’s Research UK major project grant ARUK-PG2017B−10.

## Author contributions

R.E.M., D.A.G., R.F.H., D.L.Mc.C., S. B. Z., and K. S. were responsible for the conception and design of the study. D.A.G., R.F.H., D.L.Mc.C., and S. B. Z. carried out the data analyses. R.E.M. and D.A.G. drafted the article. C.N., and A.C., facilitated data linkage. R.M.W., S.E.H., R.F., L.S., E.M.T.D., A.F.M., I.J.D., C.G., A.P., M.W., D.J.P., C.H., P.M.V., S.R.C., K.L.E., and A.M.M., contributed to data collection and preparation. All authors read and approved the final manuscript.

## Ethics

All KORA participants have given written informed consent and the study was approved by the Ethics Committee of the Bavarian Medical Association.

All components of GS received ethical approval from the NHS Tayside Committee on Medical Research Ethics (REC Reference Number: 05/S1401/89). GS has also been granted Research Tissue Bank status by the East of Scotland Research Ethics Service (REC Reference Number: 20/ES/0021), providing generic ethical approval for a wide range of uses within medical research.

Ethical approval for the LBC1921 and LBC1936 studies was obtained from the Multi-Centre Research Ethics Committee for Scotland (MREC/01/0/56) and the Lothian Research Ethics committee (LREC/1998/4/183; LREC/2003/2/29). In both studies, all participants provided written informed consent. These studies were performed in accordance with the Helsinki declaration.

## Availability of data and materials

The datasets generated and/or analysed during the current study are not publicly available.

The informed consent given by the KORA study participants does not cover posting of participant level phenotype and genotype data in public databases. However, data are available upon request from KORA-gen (http://epi.helmholtz-muenchen.de/kora-gen). Requests are submitted online and are subject to approval by the KORA board.

Lothian Birth Cohort data are available on request from the Lothian Birth Cohort Study, University of Edinburgh. Lothian Birth Cohort data are not publicly available due to them containing information that could compromise participant consent and confidentiality.

According to the terms of consent for GS and GS:STRADL participants, access to data must be reviewed by the GS Access Committee. Applications should be made to.

Code is available with open access at the following Gitlab repository: https://gitlab.com/dannigadd/episcores-for-protein-levels.

## Competing interests

R.E.M has received a speaker fee from Illumina and is an advisor to the Epigenetic Clock Development Foundation. All other authors declare no competing interests.

## Additional files

### Supplementary file 1

A. Demographic and array information for the cohorts and samples used in the study.
B. SomaScan panel performance in the STRADL test set.
C. Performance of Olink protein EpiScores in holdout, STRADL and LBC1921 test sets.
D. Annotations for the proteins corresponding to the 109 selected EpiScores.
E. Predictor weights for the 109 selected EpiScores.
F. CpG feature counts for the 109 selected EpiScores.
G. Frequency of CpG sites selected for EpiScores with EWAS catalogue annotations to phenotypic traits.
H. Basic Cox model results in Generation Scotland.
I. Fully-adjusted and sensitivity analyses results for Cox models in Generation Scotland.
J. Schoenfeld residual Cox sensitivity analyses.
K. Schoenfeld residual Cox sensitivity analyses split by year of follow-up.
L. SOMAscan – EpiScore diabetes association lookup against two large-scale plasma protein – diabetes studies.
M. White blood cell sensitivity analyses.
N. GrimAge sensitivity analyses.

### Supplementary file 2

A. Correlation structures for the 109 selected EpiScores.
B. Correlation structures for the 109 selected EpiScores in relation to common covariates.

### Supplementary file 3

A. Summary of the rationale for including each of the 12 morbidities in this study.

### Supplementary file 4

(A-L) Primary and secondary health codes for each of the 12 morbidities in this study that were used to assign case/control status of participants.

## Notes

### Competing Interest Statement

R.E.M has received payment from Illumina for presentations. All other authors declare no competing interests.

### Summary of Updates

Updates to the manuscript have been made based on review feedback to include several additional sensitivity analyses. The core results remain unchanged.

