## Supplementary file 2 for "Epigenetic scores for the circulating proteome as tools for disease prediction"

**Supplementary file 2A. Correlation heatmap for protein EpiScore measures in Generation Scotland.**

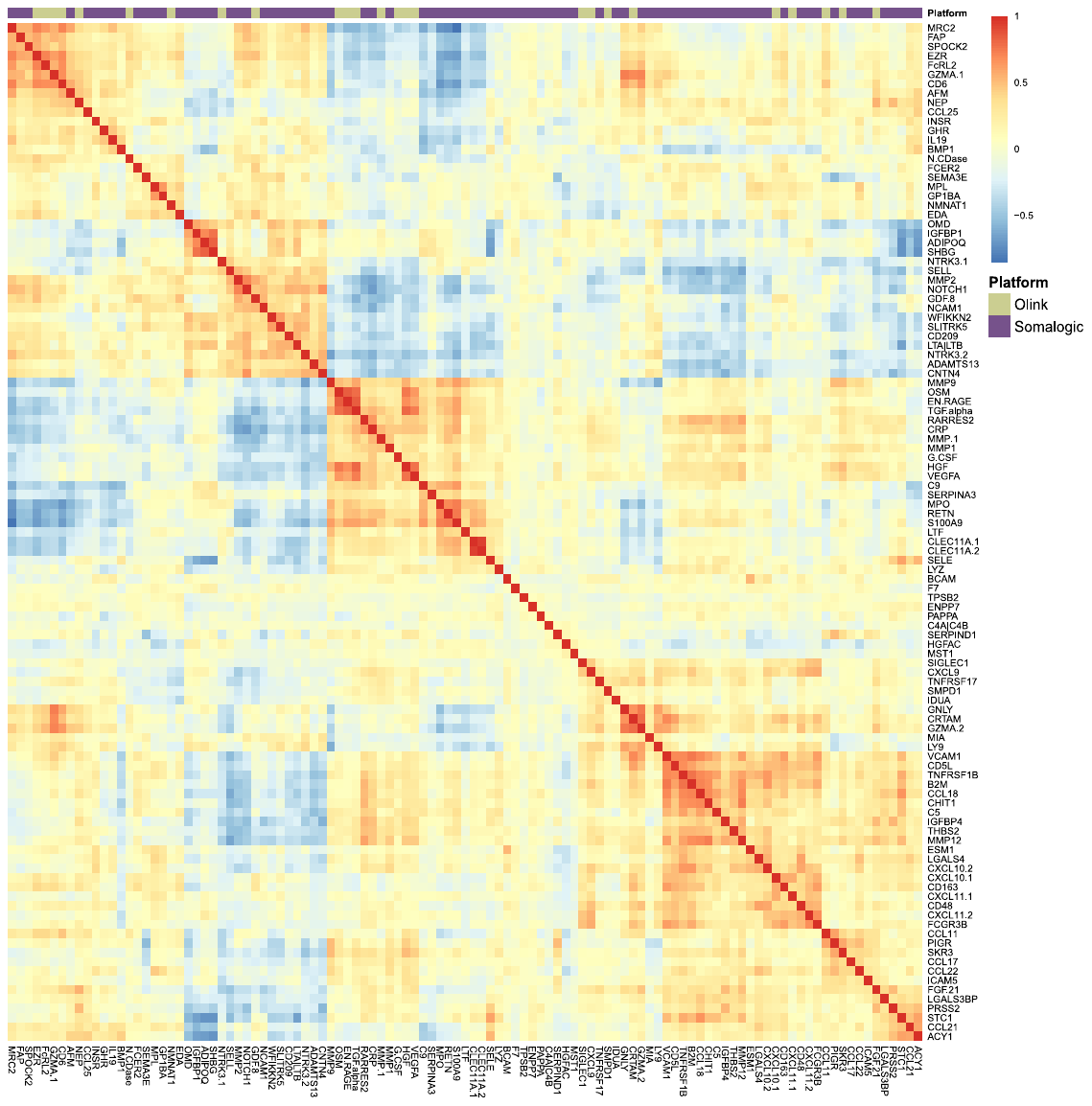

Correlation heatmap for EpiScore measures projected into Generation Scotland (N=9,537) for the 109 protein EpiScores selected in the test sample (*r* > 0.1, P < 0.05). At the top of the heatmap, an annotation bar is displayed. Olink® proteins are shown in pale green and Somalogic proteins are shown in purple.

**Supplementary file 2B. Phenotypic trait and estimated white blood cell proportion correlations with EpiScores.**

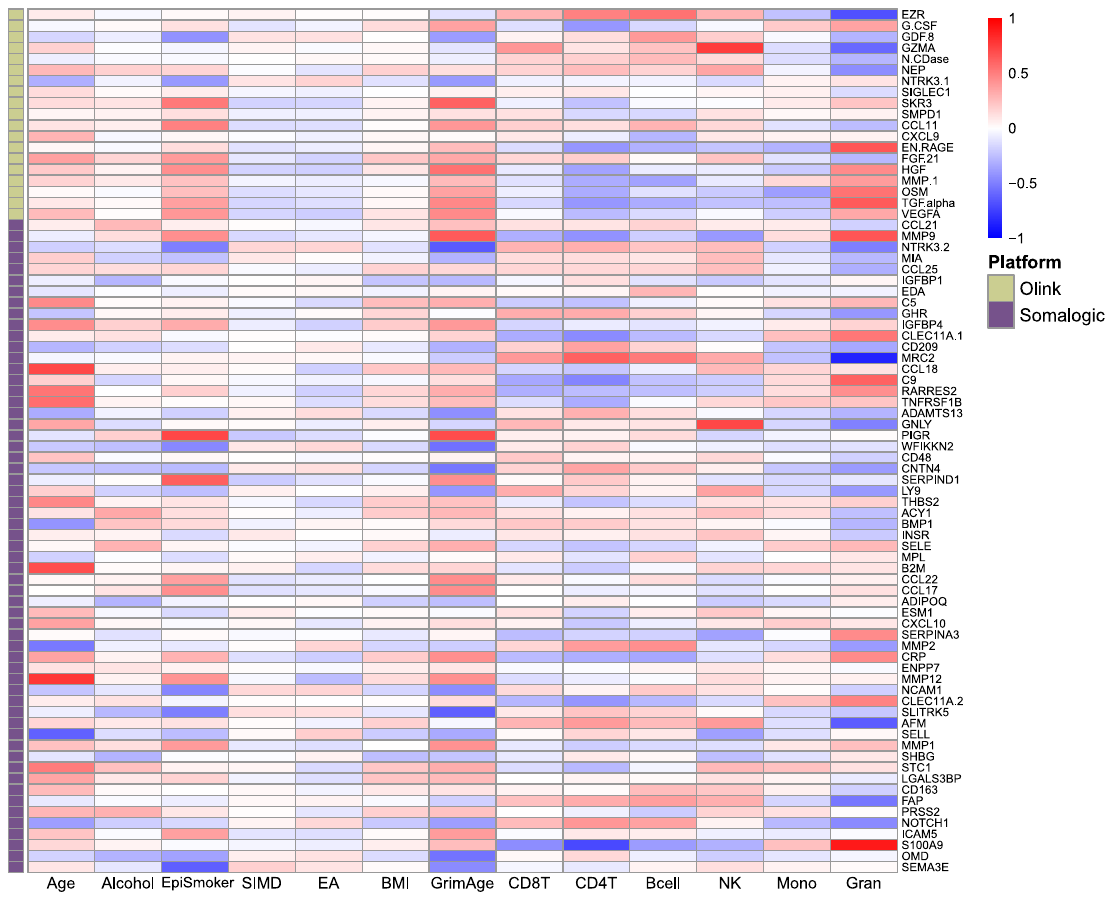

Heatmap of correlations between the 78 protein EpiScore measures for Olink® proteins which were associated with incident disease at P < 0.05 in the fully-adjusted Cox mixed effects proportional hazards models and continuous phenotypic/lifestyle trait variables (**a**) and Houseman-estimated white blood cell proportions (**b**) in Generation Scotland (total N=9,357). Protein measurements used to train the predictors were adjusted for age and sex. The maximum sample size available was used for each correlation. GrimAge: GrimAge acceleration. Units: weekly units of alcohol. EpiSmoker: DNAm-derived score for smoking. SIMD: Scottish Index of Multiple Deprivation. EA: educational attainment. Mono: Monocytes. Gran: Granulocytes. NK: Natural Killer cells.
