## Supplementary file 3 for "Epigenetic scores for the circulating proteome as tools for disease prediction"

**Supplementary file 3A: The 12 leading causes of morbidity and mortality included through data linkage.** Ten of the diseases are listed by the world health organisation (WHO) as leading causes of either morbidity or mortality. The remaining two, (rheumatoid arthritis and inflammatory bowel diseases), are also known to place morbidity burdens on individuals.

**Ten leading causes of morbidity and mortality listed by WHO**

The World Health Organisation specifies ten leading causes of mortality and ten leading causes of disease burden. In high-income countries, six diseases are present in both sets: ischemic heart disease, stroke, lung cancer, Alzheimer’s disease (AD) and other dementias, diabetes and chronic obstructive pulmonary disease (COPD). The remaining four leading causes of mortality are lower respiratory tract diseases, bowel cancer, kidney disease and breast cancer ^1^. The additional four causes of disease burden are back or neck pain, skin disease, sense organ disease and depression ^2^.

Present in WHO leading causes of mortality and causes of disease burden:

- ischemic heart disease
- stroke
- lung cancer
- Alzheimer’s disease (AD) and other dementias
- Diabetes
- chronic obstructive pulmonary disease (COPD)

Present in WHO leading causes of mortality only:

- lower respiratory tract diseases
- bowel cancer
- kidney disease
- breast cancer

Additional causes of WHO disease burden:

- back or neck pain
- skin disease
- sense organ disease
- depression

**Additional traits included in this study**

Inflammatory bowel disease (IBD) ^3^ and rheumatoid arthritis (RA) ^4^ are also leading causes of disability and morbidity and the global burdens of these diseases are rising, making them important public health issues ^5,6^.
